# Prediction of DNA i-Motifs Via Machine Learning

**DOI:** 10.1101/2023.12.11.571121

**Authors:** Bibo Yang, Dilek Guneri, Haopeng Yu, Elisé P. Wright, Wenqian Chen, Zoë A. E. Waller, Yiliang Ding

## Abstract

i-Motifs (iMs), are secondary structures formed in cytosine-rich DNA sequences and are involved in multiple functions in the genome. Although putative iM forming sequences are widely distributed in the human genome, the folding status and strength of putative iMs vary dramatically. Much previous research on iM has focused on assessing the iM folding properties using biophysical experiments. However, there are no dedicated computational tools for predicting the folding status and strength of iM structures. Here, we introduce a machine learning pipeline, iM-Seeker, to predict both folding status and structural stability of DNA iMs. The programme iM-Seeker incorporates a Balanced Random Forest classifier trained on genome-wide iMab antibody-based CUT&Tag sequencing data to predict the folding status and an Extreme Gradient Boosting regressor to estimate the folding strength according to both literature biophysical data and our in-house biophysical experiments. iM-Seeker predicts DNA iM folding status with a classification accuracy of 81% and estimates the folding strength with coefficient of determination (R^2^) of 0.642 on the test set. Model interpretation confirms that the nucleotide composition of the C-rich sequence significantly affects iM stability, with a positive correlation with sequences containing cytosine and thymine and a negative correlation with guanine and adenine.

**GRAPHICAL ABSTRACT:** 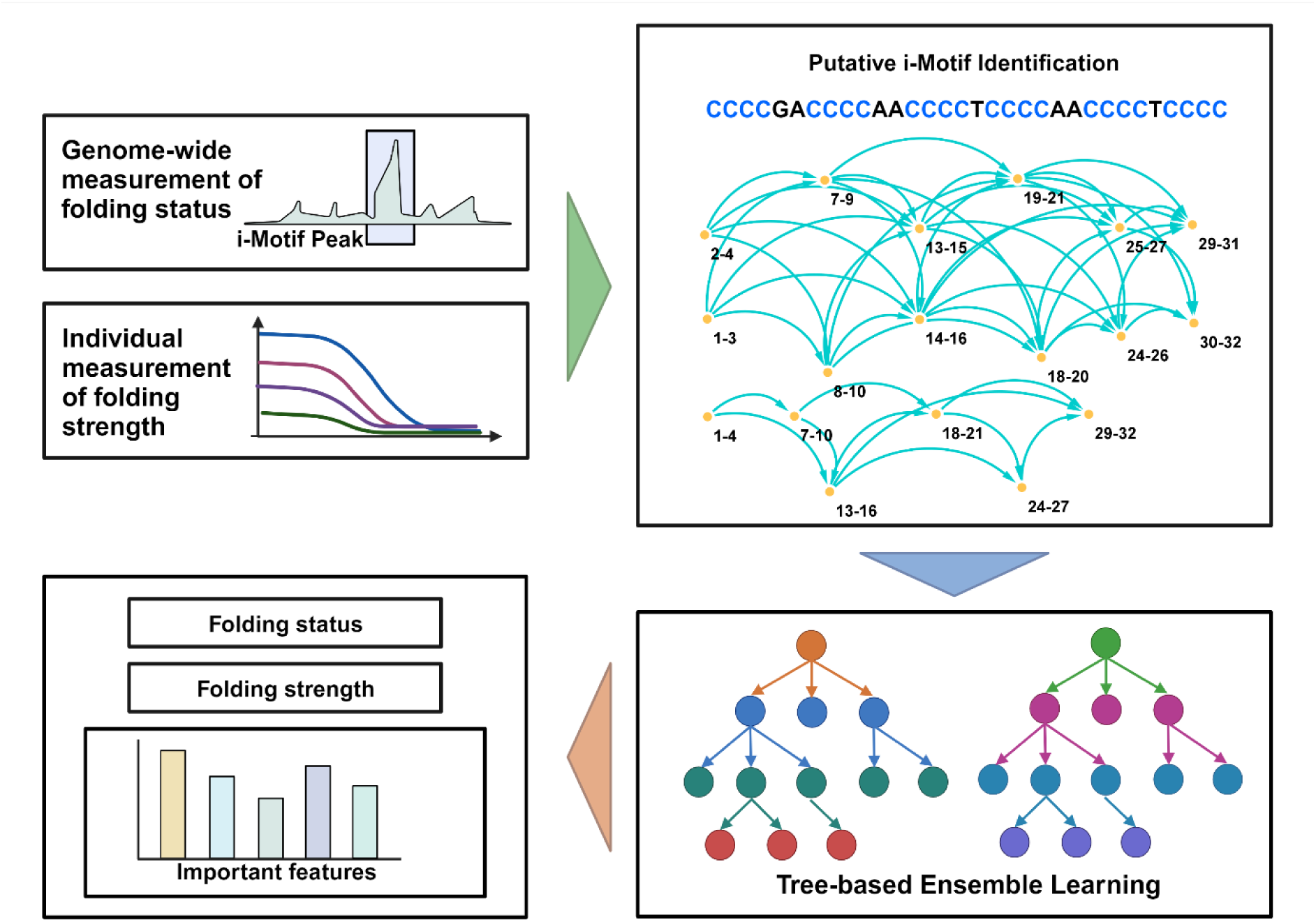

## INTRODUCTION

Nucleotides are the basic units that form DNA and RNA, two key molecules in the central dogma. DNA encodes genetic information, which is transcribed to mRNA and then translated to protein. In addition to this transfer of information, DNA and RNA can form complex structures, which can play crucial functional roles in organisms. Besides the canonical Watson-Crick double-helical B-form structure, DNA can form non-canonical secondary structures such as G-quadruplexes (G4s) and i-Motifs (iMs). G4s are four-stranded structures formed from G-rich sequences and are stabilised by Hoogsteen hydrogen bonding between guanines (1). iMs are also four-stranded structures, but formed from cytosine C-rich regions that are stabilised by hemi-protonated C-C base pairs (C^+^:C) (2,3). Complementary G-rich and C-rich sequences can form G4s and iMs interdependently during distinct cellular processes (4). As a non-canonical structure, iMs are indicated to play an important role in the genome. There are an increasing number of *in vitro* and *in celluo* studies that report evidence that iMs could fold in promotor region of certain genes, telomeres and untranslated regions. They have also been implicated as a regulatory element associated with the cell cycle, transcription, chromatin remodelling, as well as transposable element dynamics (5–7).

Commonly, computational analysis of putative iMs is limited to indirect identification by searching for potential complementary G4 sequences in the genome (8). Plenty of G4 prediction tools have been developed previously, and these can generally be divided into two categories based on whether or not the models utilised experimentally-derived G4-specific data. Classical computational tools which do not use G4-specific data, are typically constructed from string-matching models based on a specific sequence pattern. Others use a designed scoring system according to pre-defined rules. For example, platforms like Quadparser (9), Quadruplexes (10), and AllQuads (11), used algorithms like regular expression to search G4 forming sequences, whilst QGRS Mapper (12), G4P (13), and G4hunter (14), use scoring models that can estimate the probability or strength of putative G4s (15). These models have potential to be used in iM-forming sequences searching, because the putative iMs have in principle similar sequence patterns and some of the rules will be transferrable to both structures. For example, enrichment of G/C in a C/G-rich sequence disfavours both G4 and iMs. In contrast, there are also platforms guided by G4-specific data (e.g., biophysical properties, G4 ChIP-seq, G4 CUT&Tag, and G4-seq) that can capture additional G4-specific features to improve the G4 prediction performance (16). Software like PQSfinder (17), G4boost (18), Quadron (19), DeepG4 (20), and G4-folding energy estimation module integrated in RNAFold (21) use data from G4-specific experiments to increase the accuracy of predictions. Therefore, the application of these models on iM identification is limited. Out of the existing searching platforms, G4-Hunter is the easiest to use for searching for iMs as it was designed to take into account C with negative values both to disfavour regions rich in alternative G/C and to score both strand of a DNA duplex simultaneously. C-richness and C-skew is obviously important for iM formation (14). Besides, G4-iM Grinder can also be used to predict and evaluate G4 and iM forming sequences (22). Typically, individual C-rich sequences are biophysically assed for their capability to form iMs. UV spectroscopy is typically used to determine the thermodynamic properties such as melting (*T*_M_) and annealing (*T*_A_) temperatures (23). Furthermore, thermal difference spectra (TDS) are typically generated, using the difference in absorbance spectra between folded and unfolded DNA, determining a signature to identify the formed secondary DNA structure (24). UV spectroscopy is often accompanied with circular dichroism (CD) spectroscopy to conform the formation of i-motif structure. The transitional pH (pH_T_) is an important measure of the stability of iM structures, determined by assessing the formation of iM across a pH-range (8,25–27).

A systematic prediction tool to identify DNA iM folding status and their potential stability is lacking. Recently, the landscape of iM forming sequences in the whole human genome was determined via the novel CUT&Tag sequencing using anti-iM iMab antibodies on living human cells (7). Here we introduce, iM-Seeker, a novel computational pipeline using the genome-wide iM profile (7), iM-stability data from the literature, and our in-house biophysical analysis to predict iM structure formation and stability. iM-Seeker utilised a newly-designed graph-based algorithm to search for putative iM forming sequences within an entered DNA sequence. The Balanced Random Forest script is trained on the iMs identified in the human genome derived from iMab-based CUT&Tag sequencing data (7) and was further developed to predict iM structure folding status within DNA sequences. iM-Seeker also incorporates the Extreme Gradient Boosting (XGBoost) regressor to predict the structure stability, by cross referencing iM forming DNA sequences to their corresponding pH_T_ values. Furthermore, this computational model has shed new insight into the importance of nucleotide composition in iM stability. A positive correlation was observed for sequences containing cytosine and thymine whilst sequences rich in guanine and adenine were found to have a negative correlation with iM stability. Alongside nucleotide composition, long C-tract lengths accompanied with short loop lengths contribute towards high stability of iM structure.

## MATERIAL AND METHODS

### Data collection

We collected the published CUT&Tag sequencing data in the human genome (7). The data was downloaded from the NCBI GEO database (accession number GSE220882). The BigWig format data included iM forming sequences from both 93T449 (WDLPS) cell line and human embryonic kidney (HEK293T) cell line with three biological replicates for each cell line. The focus was concentrated on HEK293T cell data which was presented with more high-confident iM regions than WDLPS cells (7). The downloaded BigWig files were converted to bedGraph files and iM-peak region were cumulated with SEACR v1.3 set to “0.01 non stringent” parameters (7,28). The intersected iM-peak regions among three biological replicates were defined as the final high-confident iM-peak regions. Literature-derived data of i-Motif forming sequences and their corresponding pH_T_ values were collected (Supplementary Table S1).

### Graph-based putative i-motif searching

Putative i-Motifs can be identified based on their sequence pattern (C_≥3_N_1–12_)_3_C_≥3_ where C represents cytosine and N represent any nucleotide (29,30). The classic approach to identify potential putative iM-forming sequences is to search complementary sequences of G4-forming sequences based on sequence pattern matching. This assumption and current approaches limit the identification of iMs with their different variations in C:C(+) formations and topologies compared to G4s (29,30). To overcome this limitation, we designed a general pattern for iM formation searching using directed graph traversal process. For one sequence, the C-tracts can be regarded as nodes, and the loops can be defined as edges. All possible C-tracts (C-tract length ≥3) are identified as nodes in the first phase, and if the distance between two nodes (loop length) is between one and twelve nucleotides, a directed edge is added between the two nodes. After constructing the directed graph, all possible iM formations and conformations are identified via the traversal of the directed graph from every node. All possible putative iMs are represented with the sub-population containing the first four nodes and three edges of the traversing paths with at least four nodes. To choose the representative iM structures from all possible iM structures, four strategies were introduced (greedy non-overlapping, greedy overlapping, non-greedy non-overlapping, and non-greedy overlapping) maintaining the nomenclature derived from QuadBase2 (31). Overlapping strategy selects an iM representative structure for each iM starting coordinate while the non-overlapping function has no coinciding iM representatives. The greedy strategy maximises the loop length of iM representatives with longest C-tract. For non-greedy strategies, the iM with the most extended C-tract length and the shortest loop length can be selected. One representative iM forming sequence may have many different iM conformations although they share the same sequence content. Two representative iM formations are chosen for according to their stability: (A) the structure with minimum standard deviation of loop lengths; (B) the structure with minimum length of the two side loops. We called the initial computational pipeline Putative-iM-Searcher (Figure 1A).

**Figure 1.**
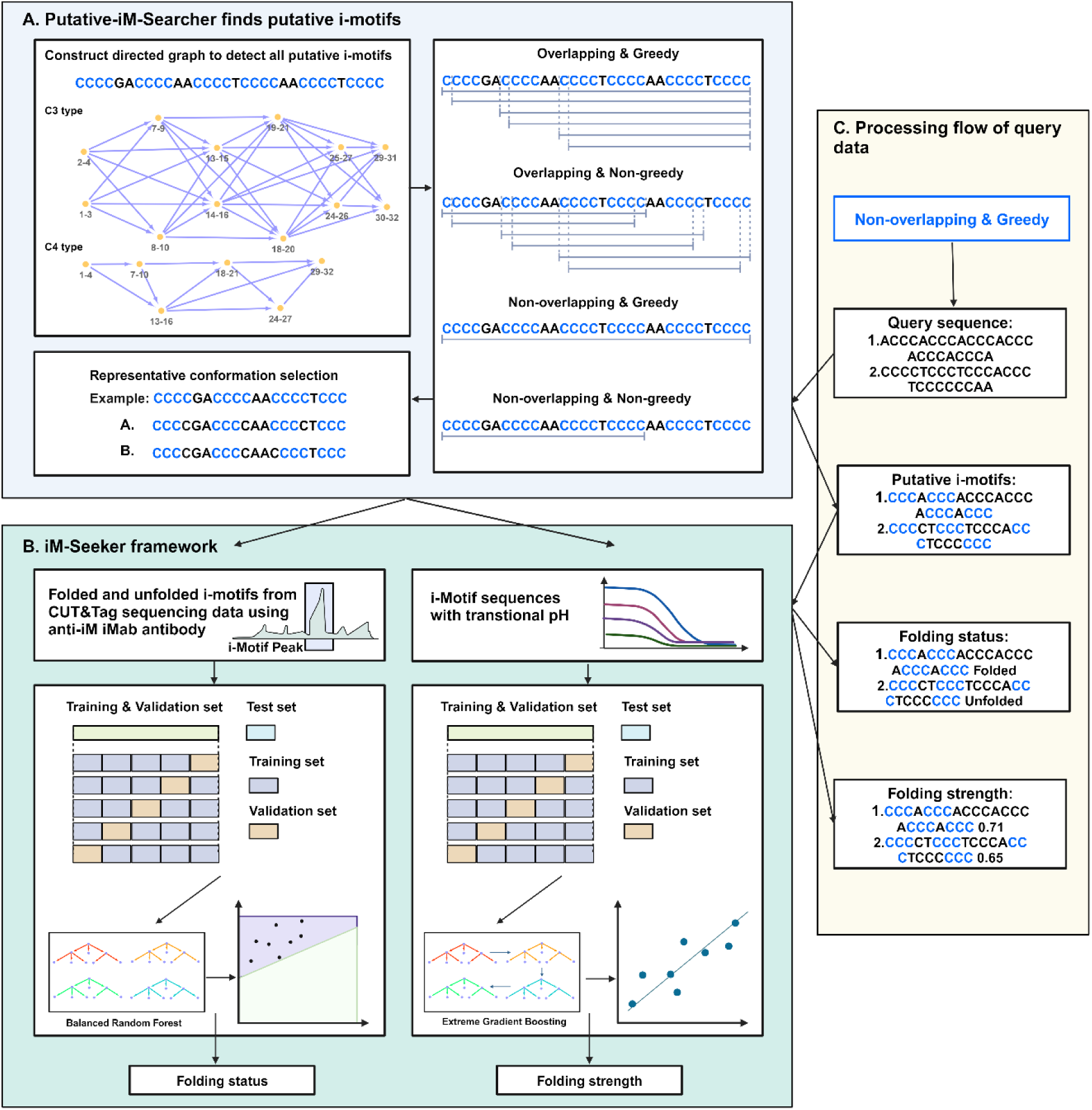
The outline of the whole iM-Seeker. (**A**) The framework of Putative-iM-Searcher. Putative-iM-Searcher can detect all i-motif conformation and representative conformation based on overlapping & non-overlapping strategy, greedy & non-greedy strategy, and representative-conformation strategy. (**B**) The framework of iM-Seeker. iM-Seeker employs Putative-iM-Searcher to find putative i-motif forming sequences. Balanced Random Forest classification model and XGBoost regression model are developed to predict the folding status and folding strength, respectively. (**C**) The processing flow of query data using iM-Seeker. Created with BioRender.com.

### Dataset construction and feature selection for machine learning

We employed a Putative-iM-Searcher in high-confident iM-peak regions and interval regions in both Watson and Crick strands in the human reference genome (GRCh38). Putative iMs in high confident iM-peak regions were defined as folded iMs, and unfolded C-rich sequences in interval regions. We used a non-overlapping strategy to avoid bias in the performance estimation of the classification model. Four classification datasets were constructed: (Classification dataset 1) non-overlapping, greedy and conformation A; (Classification dataset 2) non-overlapping, greedy and conformation B; (Classification dataset 3) non-overlapping, non-greedy and conformation A; (Classification dataset 4) non-overlapping, non-greedy and conformation B.

We selected the data items with reliable pH_T_ from literature-derived data. We also generated our in-house biophysical experimental data for developing regression models. The Putative-iM-Searcher was applied to filtered dataset of iM forming sequences with their corresponding pH_T_ values. The iMs which meet the sequence pattern with corresponding pH_T_ were used for regression model construction. We also filtered iM items with the same putative iM forming sequence but different pH_T_ and combined iM items with the same putative iM forming sequence and pH_T_ to avoid bias. Both our classification model and regression model used thirty-three different features: C-tract length, iM length, loop length, middle loop length, longest side loop length, shortest side loop length, sum of two side loops, longest loop length, shortest loop length, A density in iMs, C density in iMs, G density in iMs, T density in iMs, A density in loops, C density in loops, G density in loops, T density in loops, A density in middle loop, C density in middle loop, G density in middle loop, T density in middle loop, A density in longest side loop, C density in longest side loop, G density in longest side loop, T density in longest side loop, A density in shortest side loop, C density in shortest side loop, G density in shortest side loop, T density in shortest side loop, A density in two side loops, C density in two side loops, G density in two side loops, T density in two side loops. For the regression system, the iM folding strength is defined as the pH_T_ after standardization and min-max scaling.

### The imbalanced ensemble learning to predict folded and unfolded i-motifs

A five-fold cross-validation assessment was applied to evaluate the classification performance of the iMs for four datasets via nine classifiers including Decision Trees (32), Random Forest (33), Balanced Random Forest (34), Naive Bayes (35), Linear Discriminant Analysis (36), Easy Ensemble (37), Balanced Bagging (38,39), Random Undersampling Boosting (RUSBoost) (40), and Extreme Gradient Boosting (XGBoost) algorithms (41). The combination of dataset and model which achieve best performance via area under the receiver operating characteristic curve (AUROC) and balanced accuracy, was used for classification. 90% of data in the whole dataset was randomly selected and separated into a training & validation set, and the remaining 10% of data was used as the test set. Five-fold cross-validation and grid searching on training & validation set were employed to search for the best hyperparameters and test set was used to evaluate the model’s classification performance on accuracy, recall, specificity, and AUROC.

### The regression algorithm to measure the strength of i-motif using ensemble learning

Consistent iM searching and conformation identification strategy with classification dataset was applied in the regression model. A five-fold cross validation assessment was applied to evaluate the regression performance of the iMs based on thirteen regressors including Decision Trees (32), Random Forest (33), Linear Regression (42), Ridge Regression (43), Lasso Regression (44), Elastic Net Linear Regression (45), Linear Support Vector Regression (46), Radial Basis Function Support Vector Regression (47), K-Nearest Neighbors Regression (KNN) (48), Adaptive Boosting (AdaBoost) (49), Gradient Boosting (50), Extreme Gradient Boosting (XGBoost) (41), and Random Sample Consensus (RANSAC) algorithms (51). 80% of data in the whole dataset was separated into training & validation set randomly for hyperparameters adjustment by five-fold cross-validation and grid searching, and 20% of data was used to evaluate the regression performance of the model by coefficient of determination (R^2^), root mean squared error (RMSE), and mean absolute error (MAE) (18). The feature importance of the regression model was extracted from the model with ‘importance_type=gain’.

### Implementation

The algorithm was written in Python 3, and machine learning was employed via the Python Scikit-learn package (52), Imbalanced-learn package (53), and XGBoost package (41). The source code and documentation of Putative-iM-Searcher are available at https://github.com/YANGB1/Putative-iM-Searcher. Combining the classification model and regression model, we built a computational tool called iM-Seeker, which is available at https://github.com/YANGB1/iM-Seeker.

### Biophysical Characterisation of C-rich DNA sequences

The test oligonucleotides were synthesised and reverse phase HPLC purified by Eurogentec (Belgium) and were resuspended in ultra-pure water. The DNA final concentration was confirmed via Nanodrop. Samples were prepared as 10 µM DNA in 10 mM sodium cacodylate (NaCaco) and 100 mM KCl buffer with the range of pH 4-8. The DNA samples were annealed prior to biophysical characterisation by denaturing the DNA for 5 mins at 95°C and allowing to reanneal by slowly cooling down to room temperature, overnight.

The CD spectra of the annealed C-rich sequences were recorded on a JASCO 1500 spectropolarimeter under a constant flow of nitrogen. An accumulation of four CD spectra scans was acquired from 200-320 nm at 20°C with a data pitch of 0.5 nm, scanning speed of 200 nm/min with 1 second response time, 1 nm bandwidth, and 200 mdeg sensitivity. The measured DNA samples and buffer at corresponding pH were subtracted before zero correction at 320 nm. The transitional pH (pH_T_) was determined by plotting the measured ellipticity at 288 nm and pH range and the resulting inflection point of the Boltzmann sigmoidal or bi-phase sigmoidal fit using Graphpad Prism (Version 10.1.0.316).

The CD samples at pH 5.5 were diluted in the same buffer to 2.5 µM final DNA concentration. These samples were used to perform UV spectroscopy to obtain the thermal difference spectra (TDS) and determine the melting temperature (T_M_), annealing temperature (T_A_) and their respective hysteresis (T_H_). For melting/annealing experiments, the absorbance at 295 nm was measured at every 1°C increase/decrease in three cycles of denaturation and reannealing. The cycle begins with 10 mins at 4°C followed by gradual increase of 0.5°C/min to 95°C (melting). Once the final temperature was reached, the samples were kept at 95°C for 10 mins before reversing the process (annealing). The melting and annealing temperatures were determined via the first derivative method of for each measured cycle as previously

described (54). The samples were kept at 4°C after the completion of the final reannealing cycle. For the thermal difference spectra (TDS), these samples were used to obtain the absorbance spectrum (230-320 nm). The samples were kept at 4°C for an additional 10 mins before measuring the absorbance spectrum of potentially folded iMs. This was followed by a second absorbance spectrum after 10 mins at 95°C for the unstructured DNA structure. Individual TDS signatures were determined by subtracting both absorbance spectra (unfolded-folded DNA structure), zero correcting at 320 nm, and finally normalisation to the maximum absorbance to 1 as previously described (24).

## RESULTS

### Description of the iM-Seeker framework

iM-Seeker is a computational framework using machine learning to predict the folding status and folding strength of iMs. The outline of the whole iM-Seeker structure is shown in Figure 1. The Putative-iM-Searcher was developed to discover the putative iM forming sequences (Figure 1A). Putative-iM-Searcher constructs a directed graph model and obtains representative conformation from all DNA structure conformations based on the configuration of overlapping & non-overlapping strategy, greedy & non-greedy strategy, and representative-conformation-selection strategy. The Balanced Random Forest classification model and XGBoost regression model were trained on iMab-based genome-wide iM landscape and biophysical experimental justified iM with pH_T_, respectively, for the folding status prediction and folding strength estimation (Figure 1B). The workflow of iM-Seeker after receiving the query sequences is shown in Figure 1C. Putative-iM-Searcher was applied to query sequences to find putative iM forming sequences in the first stage. For each putative iM individual, the Balanced Random Forest classification model will be used to predict the folding status. Next, an estimated folding strength score was calculated by the XGBoost regression model for putative iM individuals.

### iM-Seeker predicts iM structure folding status

Putative iM forming sequences in the intersected high-confident iM-peak regions among three biological replicates from the CUT&Tag sequencing data were defined as folded iMs while unfolded C-rich sequences can be found in interval regions. For greedy and non-overlapped two classification datasets, they both included 8,837 folded iMs and 733,115 unfolded C-rich sequences while 9,641 folded iMs and 755,747 unfolded C-rich sequences were in non-greedy and non-overlapped two datasets.

Thirty-three features from labelled folded and unfolded putative iM sequences were derived. A five-fold cross-validation assessment was applied on nine classifiers on four classification datasets to select the best dataset and model. Considering the mean AUROC score and mean balanced accuracy of five folds, Balanced Random Forest performed best in all four datasets because the balanced learning strategy can better fit our imbalanced datasets. Thus, Balanced Random Forest was selected as the final classifier. Greedy and non-overlapped two datasets outperformed the non-greedy and non-overlapped datasets in terms of the two indicators. Although there is no significant difference between conformation A and B for greedy and non-overlapped datasets, both AUROC and balanced accuracy of conformation A were found to be higher than B (Figure 2A). Thus, we chose conformation A dataset of greedy and non-overlapped strategy as final dataset for classification task.

**Figure 2.**
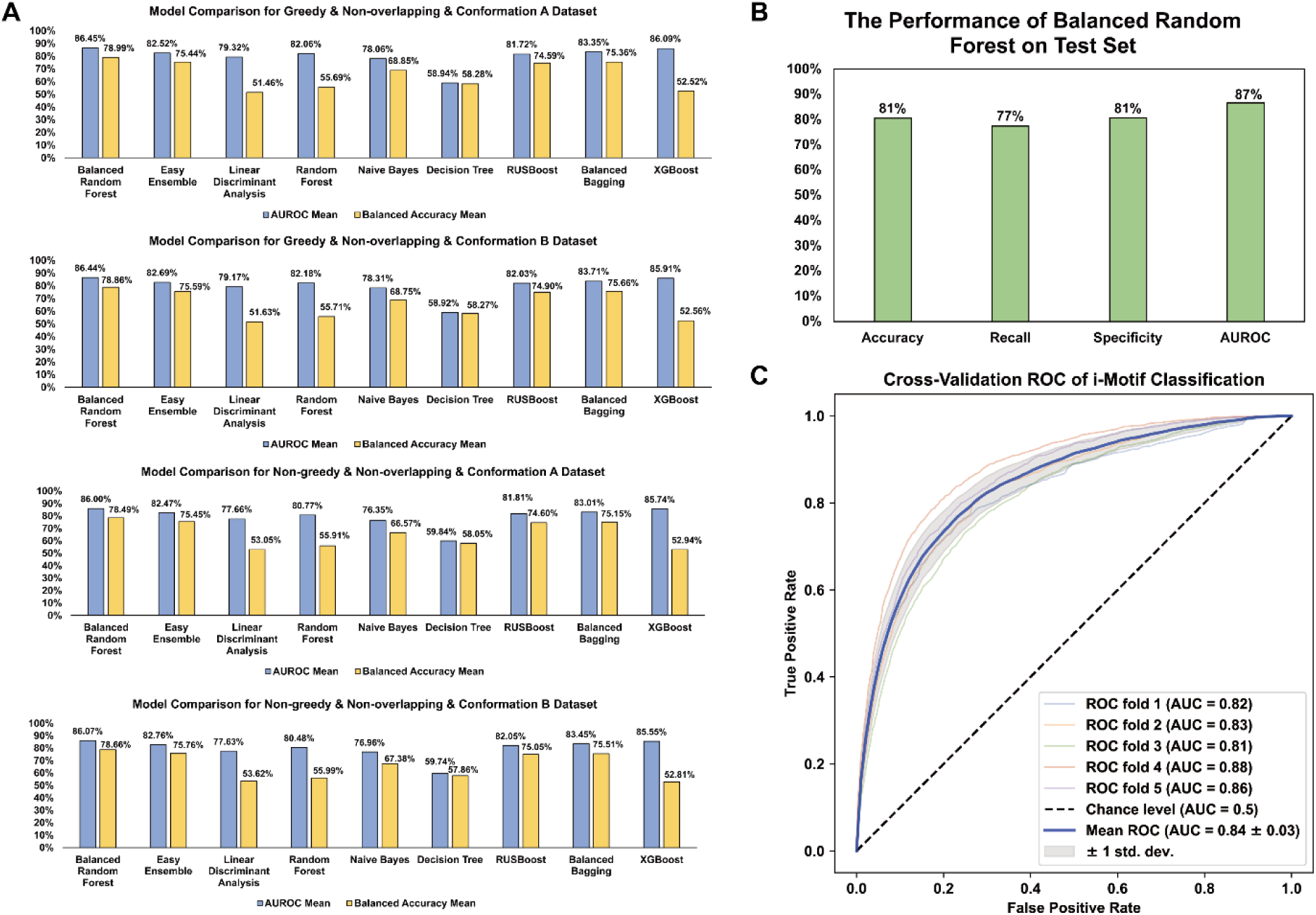
Model selection and performance estimation of classification model. (**A**) The comparison among nine models (Decision Trees, Random Forest, Balanced Random Forest, Naive Bayes, Linear Discriminant Analysis, Easy Ensemble, Balanced Bagging, RUSBoost, and XGBoost) on four classification datasets. AUROC and balanced accuracy show that Balanced Random Forest on greedy & non-overlapping & conformation B dataset has the best performance. (**B**) The performance of Balanced Random Forest classifier on the test set. Accuracy, recall, specificity, and AUROC can reach 81%, 77%, 81%, and 87% respectively. (**C**) The ROC curves for classification performance. The Receiver Operating Characteristic (ROC) for the five-fold cross validation is shown. Each fold coloured separately with the AUC score and the mean ROC curve are coloured blue, and the random probability is shown as black dash lines.

The whole dataset was divided into the training & validation set (90%) and test set (10%) because the whole dataset contains ∼740,000 data items, test set with ∼74,000 (10%) data items is enough to test the model performance. The Balanced Random Forest model was optimised by cross-validation and grid search on training & validation set. We evaluated the model performance on the test set with 81% accuracy, 77% recall, 81% specificity, and 87% AUROC score, which show the model can achieve good performance in both folded iMs and unfolded C-rich sequences (Figure 2B). Besides, we assessed the model’s generalisation performance through five-fold cross-validation deployed across the entire dataset on AUROC (Figure 2C). The AUROC scores on all five folds are all higher than 0.8, which shows the excellent generalisation performance.

### iM-Seeker measures the iM structure stability

The literature-derived data (Supplementary Table S1) and the experimental biophysical data (Supplementary Table S2) were combined to a collection of 206 C-rich DNA sequences with their corresponding pH_T_ values. The comparison of CD spectroscopy, UV spectroscopy, and TDS between representative iM forming sequence and representative non-iM forming sequence shows the reliability of our experiments (Supplementary Figure S1). However, one study contained 196 different sequences which contained only C and T. To avoid bias, these DNA sequences were excluded to avoid misinterpretation of the importance of different nucleotides in the loops. 171 data items were selected as high-confident iM-containing items from 206 items based on criteria including TDS (Supplementary Data Set 1). After filtering data items with the same putative iM but different pH_T_ and combining iM items with the same putative iM and equal pH_T_ from high-confident data items, 120 putative iMs were extracted from the remaining sequence segments using the consistent Putative-iM-Searcher strategies (greedy, non-overlapped, and conformation A) with classification session followed by feature selection (Supplementary Data Set 2). The 120 pH_T_ values standardized and rescaled to range from 0 to 1 via min-max scaling to define iM folding stability.

A five-fold cross-validation assessment was applied to thirteen regressors on regression datasets to find the model. Considering the mean of three indicators (R^2^, RMSE, and MAE) on five-folds, XGBoost was selected as the final model because of the best performance (Table 1). The whole dataset was divided into training & validation set with 80% data and test set containing the remaining data. After optimization using cross-validation and grid search on training & validation set, the final XGBoost model was applied to the test set to assess the performance. R^2^, RMSE, and MAE can reach 0.642, 0.104, and 0.08, respectively, which shows the model can achieve good performance in estimating the folding strength (Figure 3). The Pearson Correlation Coefficient (PCC) also reveals a strong correlation between measured and predicted folding strength (p<10^-7^).Model interpretation provides insights into important features for iM stability

**Figure 3.**
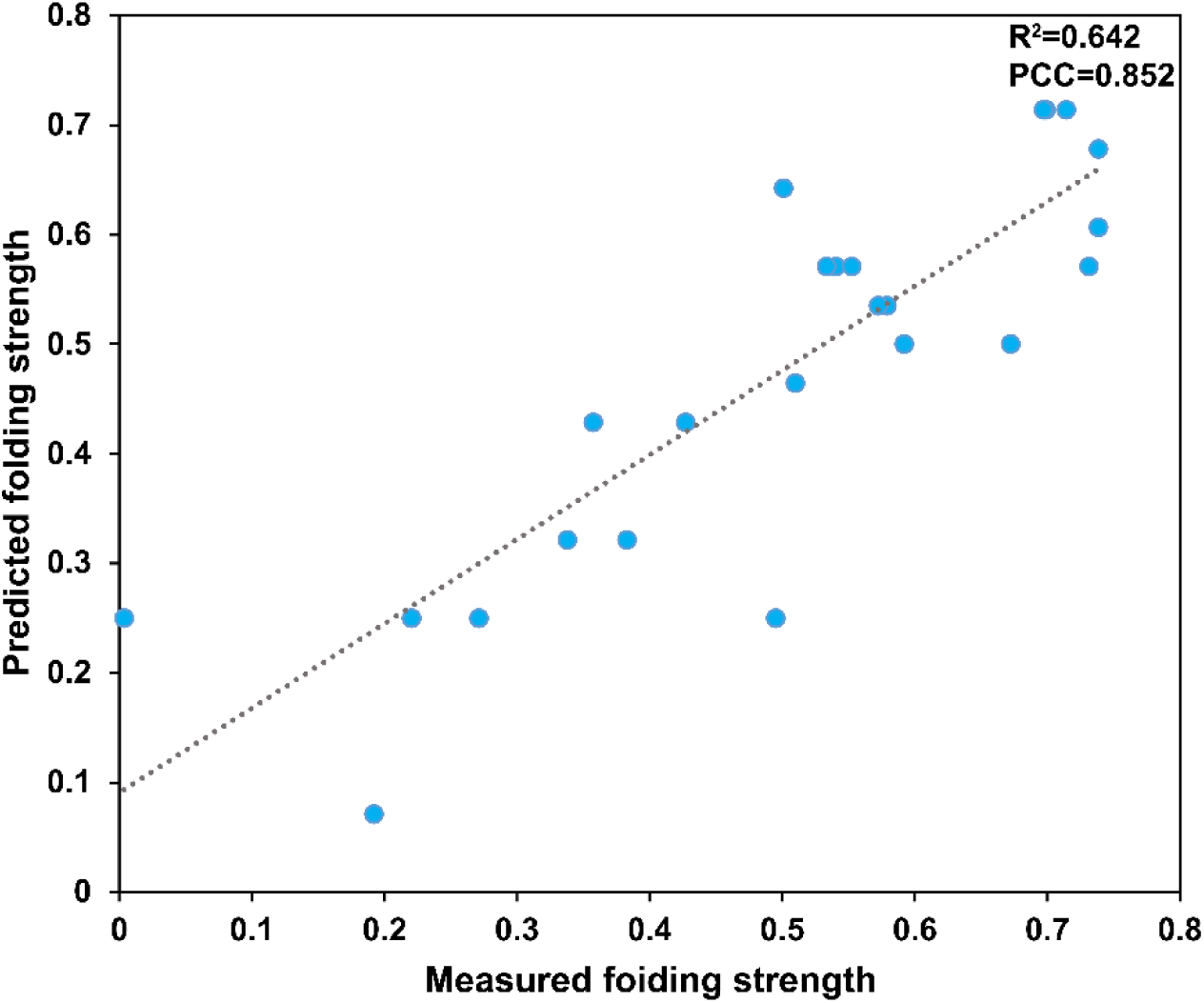
The performance evaluation of the XGBoost regressor on the test set (n=24). The Pearson Correlation Coefficient (PCC, 0.852, p<10^-7^) and R^2^ (0.642) show a positive correlation between measured and predicted iM folding strength.

**Table 1.**
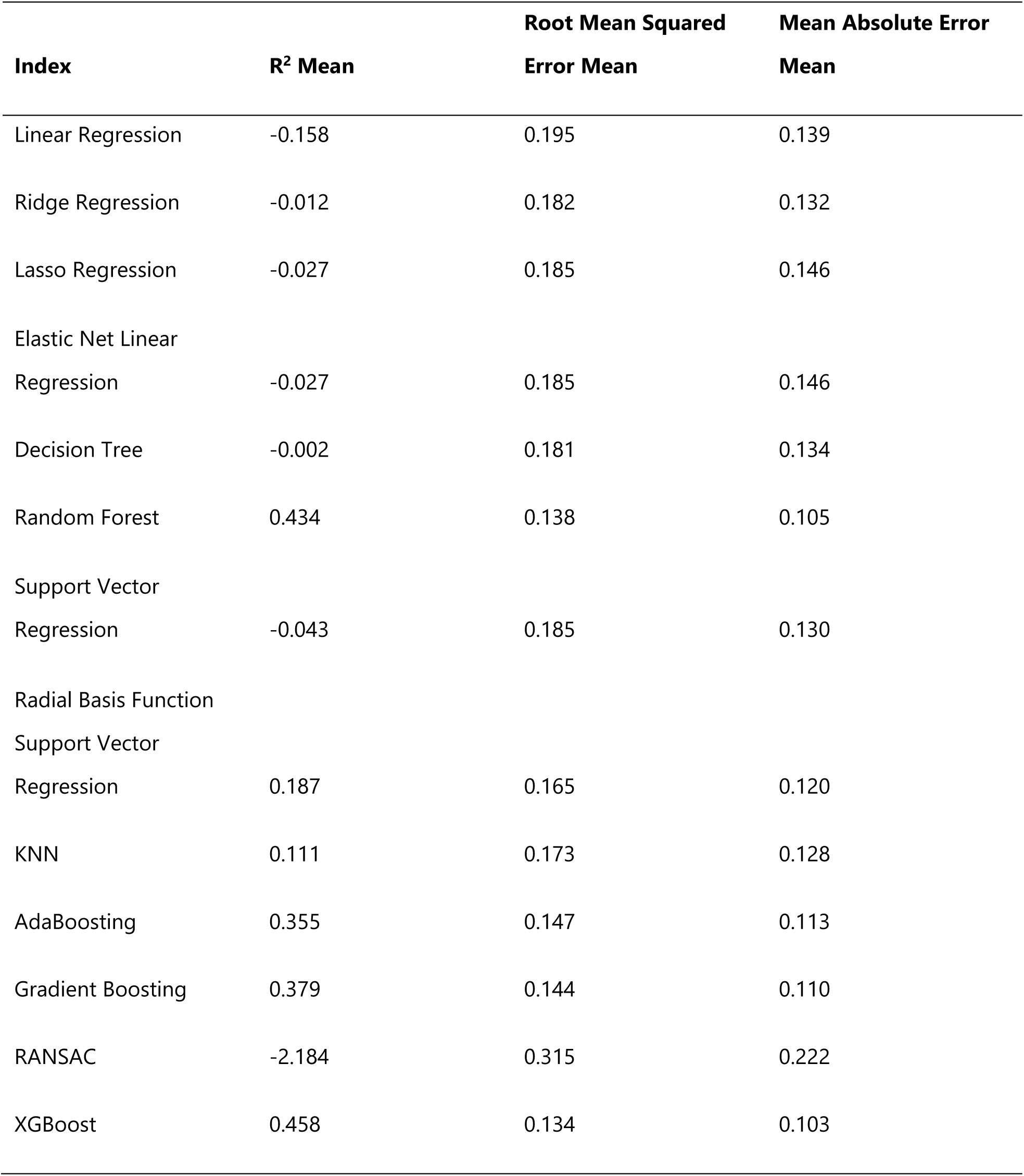
Model comparison of thirteen regressors.

| Index | R<sup>2</sup> Mean | Root Mean Squared<br>Error Mean | Mean Absolute Error<br>Mean |
| --- | --- | --- | --- |
| Linear Regression | -0.158 | 0.195 | 0.139 |
| Ridge Regression | -0.012 | 0.182 | 0.132 |
| Lasso Regression | -0.027 | 0.185 | 0.146 |
| Elastic Net Linear<br>Regression | -0.027 | 0.185 | 0.146 |
| Decision Tree | -0.002 | 0.181 | 0.134 |
| Random Forest | 0.434 | 0.138 | 0.105 |
| Support Vector<br>Regression | -0.043 | 0.185 | 0.130 |
| Radial Basis Function<br>Support Vector<br>Regression | 0.187 | 0.165 | 0.120 |
| KNN | 0.111 | 0.173 | 0.128 |
| AdaBoosting | 0.355 | 0.147 | 0.113 |
| Gradient Boosting | 0.379 | 0.144 | 0.110 |
| RANSAC | -2.184 | 0.315 | 0.222 |
| XGBoost | 0.458 | 0.134 | 0.103 |

We investigated the relative importance of the iM features extracted from the regression model. Features with high importance contribute more to the construction of the model and may play a more crucial role in iM formation than features with low importance. We divided the features into two groups based on the Pearson Correlation Coefficient (PCC): features with positive PCC were assumed to strengthen iM formation (Supplementary Table S3). In contrast, negative-correlated features were supposed to have a negative effect (Supplementary Table S4). In each group, the top 10 critical features are shown in Figure 4. Nucleotide composition affects the stability of iM structures. Stable iMs prefer to contain more C and T, especially T in side loops (Figure 4A). High G density and A density are associated with unstable iMs, especially these two nucleotides in side loops (Figure 4B). In addition, the C-tract length and loop length are two dominant features in all length-relative features. Long C-tract and short loop length can help with iM stability. Previous studies showed that in the same experimental condition, iMs with long C-tracts tend to be more stable than iMs with short C-tracts (55,56).

**Figure 4.**
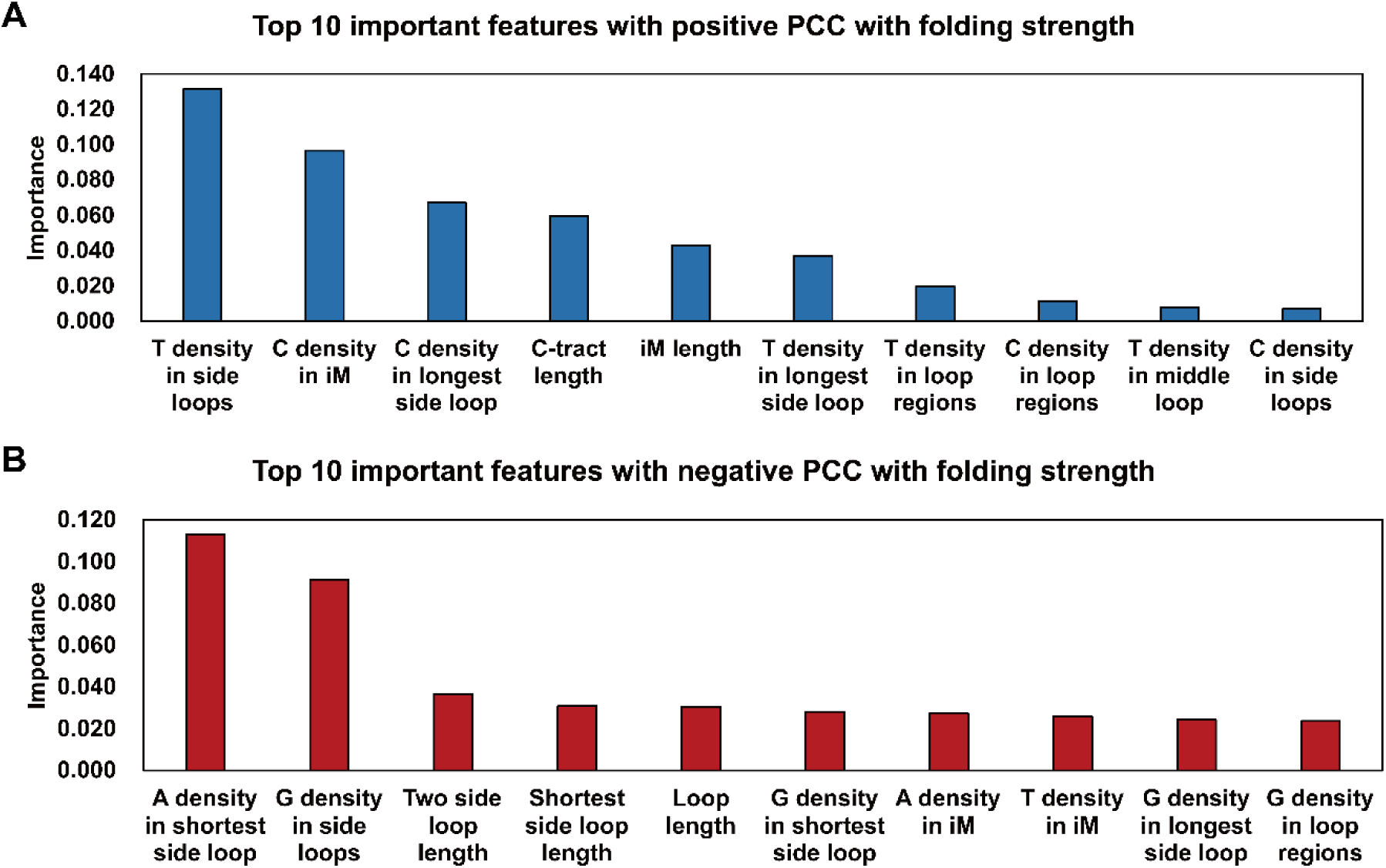
The iM feature importance obtained from the regression model. (**A**) Top 10 important features with positive Pearson Correlation Coefficient (PCC) with folding stability. (**B**) Top 10 important features with negative Pearson Correlation Coefficient (PCC) with folding stability.

## DISCUSSION

Unlike the computational prediction of G4 structures, iMs are more complex in terms of what makes them stable (8,27,29,57–60) and it has been difficult to make predictions about iMs in the same way as G4s. Although, putative iMs have a similar sequence pattern to G4s, the stability of the structures has been more difficult to predict, as it has been shown that iMs can tolerate changes in sequence more than G4s (29), but are overall less stable in general. Therefore, iM-specific experimental data is critical to construct accurate computational models for iM prediction and stability. To the best of our knowledge, there are no iM-specific computational tools. Due to the similarity in sequence patterns between G4 and iM, some previous software developed for G4 can be used on putative iM searching and can calculate a numeric value to estimate iM (8,14,22) but there was no iM-specific experimental results which were fed into models to help with model design and training. In this paper, we developed both a putative iM-forming sequence searching tool, Putative-iM-Searcher, and a machine learning approach to prediction of DNA iM folding status and folding strength, iM-Seeker. We considered that the identification of putative iM forming sequences, their folding status and folding strength were three significant parts of iM investigation that could benefit from computational predictions. Putative-iM-Searcher can construct directed graphs based on different configurations, can search all putative iM formations and conformations by graph traversal from input DNA sequences. Users can choose to set parameters including C-tract length, the first loop length, the second loop length, and the third loop length. The representative conformations can be obtained based on overlapping & non-overlapping strategy, greedy & non-greedy strategy, and representative-conformation strategy. Users can choose to obtain all putative iM formations and conformations as well. Based on the detected putative iMs by Putative-iM-Searcher, we used genome-wide CUT&Tag sequencing data and experimental data with pH_T_ from previous studies and our experiments to develop iM-Seeker. This is the first time a machine learning approach has been applied to classification of this specific DNA structure motif and will significantly improve the accuracy of *in silco* iM prediction. The iMab antibody-based CUT&Tag sequencing data presents the folding status of C-rich sequences and iM-Seeker captures the difference between features in both folded iMs and unfolded C-rich sequences and allows for classification. Another regression model was trained on iM sequences derived from biophysical data, corresponding sequence with pH_T_ to measure the folding strength.

iM-Seeker has good performance on both classification and regression tasks. The Balanced Random Forest classifier has higher performance in the imbalanced dataset. The number of folded iMs (8,837 iMs) is much less than unfolded motifs (733,115 iMs), which can mislead the classifier to overfit the unfolded dataset and classify folded iMs into unfolded category incorrectly. However, Balanced Random Forest is a decision-tree-based ensemble learning model that employs an under-sampling strategy to avoid overfitting of unfolded samples. Therefore, both folded samples and unfolded samples have good performance (recall 77%; specificity 81%). XGBoost, another ensemble learning approach which was also used in the G4 classification mission (18), is selected for the estimation of folding strength among thirteen regressors. Although the number of data items for the regression model is limited, the regression part of iM-Seeker can also provide a reliable reference to evaluate the iM strength (R^2^ 0.642; RMSE 0.104; MAE 0.08). Previous studies investigated the iM formation features which can influence the iM strength by biophysical characterisation. The length of C-tracts, short loop length and high density of C and T can enhance the formation of iMs because other strong structures can be formed with G and A, which can result in the competition between iM and other structure motifs (8,55,56,61,62). Important features extracted from the regression model revealed a consistent result with previous research, which also justifies the reliability of our model. However, the stabilising effect of additional thymines is now quite well documented and consistent with the results observed here (29,63). Also the competition between guanines and cytosines were previously used in G4-hunter (14) as a scoring factor as having the complementary base within the sequences can skew structure formation towards hairpin (29).

iM-Seeker offers users the opportunity for a dedicated iM-searching tool, which is based on machine learning from existing datasets. The approach could be applied to other DNA and RNA structures where there is a wide range of data available, for example to further increase the accuracy of prediction of formation of G4 structures.

## SUPPLEMENTARY DATA

Supplementary Data are available at NAR online.

## ACKNOWLEDGEMENT

This research was partly supported by the Norwich Bioscience Institutes Partnership’s Computing infrastructure for Science (CiS) group through the provision of a High-Performance Computing Cluster and the John Innes Centre Informatics team.

## REFERENCES

1. Lane, A.N., Chaires, J.B., Gray, R.D. and Trent, J.O. (2008) Stability and kinetics of G-quadruplex structures. Nucleic acids research, 36, 5482–5515.

2. Gehring, K., Leroy, J.-L. and Guéron, M. (1993) A tetrameric DNA structure with protonated cytosine-cytosine base pairs. Nature, 363, 561–565.

3. Kang, C., Berger, I., Lockshin, C., Ratliff, R., Moyzis, R. and Rich, A. (1994) Crystal structure of intercalated four-stranded d (C3T) at 1.4 A resolution. Proceedings of the National Academy of Sciences, 91, 11636–11640.

4. King, J.J., Irving, K.L., Evans, C.W., Chikhale, R.V., Becker, R., Morris, C.J., Peña Martinez, C.D., Schofield, P., Christ, D. and Hurley, L.H. (2020) DNA G-quadruplex and i-motif structure formation is interdependent in human cells. Journal of the American Chemical Society, 142, 20600–20604.

5. Zeraati, M., Langley, D.B., Schofield, P., Moye, A.L., Rouet, R., Hughes, W.E., Bryan, T.M., Dinger, M.E. and Christ, D. (2018) I-motif DNA structures are formed in the nuclei of human cells. Nature chemistry, 10, 631–637.

6. Ma, X., Feng, Y., Yang, Y., Li, X., Shi, Y., Tao, S., Cheng, X., Huang, J., Wang, X.-e. and Chen, C. (2022) Genome-wide characterization of i-motifs and their potential roles in the stability and evolution of transposable elements in rice. Nucleic Acids Research, 50, 3226–3238.

7. Zanin, I., Ruggiero, E., Nicoletto, G., Lago, S., Maurizio, I., Gallina, I. and Richter, S.N. (2023) Genome-wide mapping of i-motifs reveals their association with transcription regulation in live human cells. Nucleic Acids Research, 51, 8309–8321.

8. Wright, E.P., Huppert, J.L. and Waller, Z.A. (2017) Identification of multiple genomic DNA sequences which form i-motif structures at neutral pH. Nucleic acids research, 45, 2951–2959.

9. Huppert, J.L. and Balasubramanian, S. (2005) Prevalence of quadruplexes in the human genome. Nucleic acids research, 33, 2908–2916.

10. Todd, A.K., Johnston, M. and Neidle, S. (2005) Highly prevalent putative quadruplex sequence motifs in human DNA. Nucleic acids research, 33, 2901–2907.

11. Kudlicki, A.S. (2016) G-quadruplexes involving both strands of genomic DNA are highly abundant and colocalize with functional sites in the human genome. PLoS One, 11, e0146174.

12. Kikin, O., D’Antonio, L. and Bagga, P.S. (2006) QGRS Mapper: a web-based server for predicting G-quadruplexes in nucleotide sequences. Nucleic acids research, 34, W676–W682.

13. Eddy, J. and Maizels, N. (2006) Gene function correlates with potential for G4 DNA formation in the human genome. Nucleic acids research, 34, 3887–3896.

14. Bedrat, A., Lacroix, L. and Mergny, J.-L. (2016) Re-evaluation of G-quadruplex propensity with G4Hunter. Nucleic acids research, 44, 1746–1759.

15. Puig Lombardi, E. and Londoño-Vallejo, A. (2020) A guide to computational methods for G-quadruplex prediction. Nucleic acids research, 48, 1–15.

16. Elimelech-Zohar, K. and Orenstein, Y. (2023) An overview on nucleic-acid G-quadruplex prediction: from rule-based methods to deep neural networks. Briefings in Bioinformatics, 24, bbad252.

17. Hon, J., Martínek, T., Zendulka, J. and Lexa, M. (2017) pqsfinder: an exhaustive and imperfection-tolerant search tool for potential quadruplex-forming sequences in R. Bioinformatics, 33, 3373–3379.

18. Cagirici, H.B., Budak, H. and Sen, T.Z. (2022) G4Boost: a machine learning-based tool for quadruplex identification and stability prediction. BMC bioinformatics, 23, 1–18.

19. Sahakyan, A.B., Chambers, V.S., Marsico, G., Santner, T., Di Antonio, M. and Balasubramanian, S. (2017) Machine learning model for sequence-driven DNA G-quadruplex formation. Scientific reports, 7, 14535.

20. Rocher, V., Genais, M., Nassereddine, E. and Mourad, R. (2021) DeepG4: a deep learning approach to predict cell-type specific active G-quadruplex regions. PLOS Computational Biology, 17, e1009308.

21. Lorenz, R., Bernhart, S.H., Höner zu Siederdissen, C., Tafer, H., Flamm, C., Stadler, P.F. and Hofacker, I.L. (2011) ViennaRNA Package 2.0. Algorithms for molecular biology, 6, 1–14.

22. Belmonte-Reche, E. and Morales, J.C. (2020) G4-iM Grinder: when size and frequency matter. G-Quadruplex, i-Motif and higher order structure search and analysis tool. NAR genomics and bioinformatics, 2, lqz005.

23. Mergny, J.L. and Lacroix, L. (2009) UV melting of G-quadruplexes. Current protocols in nucleic acid chemistry, 37, 17.11. 11-17.11. 15.

24. Mergny, J.-L., Li, J., Lacroix, L., Amrane, S. and Chaires, J.B. (2005) Thermal difference spectra: a specific signature for nucleic acid structures. Nucleic acids research, 33, e138–e138.

25. Iaccarino, N., Di Porzio, A., Amato, J., Pagano, B., Brancaccio, D., Novellino, E., Leardi, R. and Randazzo, A. (2019) Assessing the influence of pH and cationic strength on i-motif DNA structure. Analytical and bioanalytical chemistry, 411, 7473–7479.

26. Nguyen, T., Fraire, C. and Sheardy, R.D. (2017) Linking pH, temperature, and K+ concentration for DNA i-Motif formation. The Journal of Physical Chemistry B, 121, 7872–7877.

27. Gurung, S.P., Schwarz, C., Hall, J.P., Cardin, C.J. and Brazier, J.A. (2015) The importance of loop length on the stability of i-motif structures. Chemical Communications, 51, 5630–5632.

28. Meers, M.P., Tenenbaum, D. and Henikoff, S. (2019) Peak calling by Sparse Enrichment Analysis for CUT&RUN chromatin profiling. Epigenetics & chromatin, 12, 1–11.

29. Guneri, D., Alexandrou, E., El Omari, K., Dvorakova, Z., Chikhale, R.V., Pike, D., Waudby, C.A., Morris, C.J., Haider, S. and Parkinson, G.N. (2023) Structural Insights into Regulation of Insulin Expression Involving i-Motif DNA Structures in the Insulin-Linked Polymorphic Region. bioRxiv, 2023.2006. 2001.543149.

30. Williams, S.L., Casas-Delucchi, C.S., Raguseo, F., Guneri, D., Li, Y., Minamino, M., Fletcher, E.E., Yeeles, J.T., Keyser, U.F., Waller, Z.A. et al. (2023) Replication-induced DNA secondary structures drive fork uncoupling and breakage. The EMBO Journal, 42, e114334.

31. Dhapola, P. and Chowdhury, S. (2016) QuadBase2: web server for multiplexed guanine quadruplex mining and visualization. Nucleic acids research, 44, W277–W283.

32. Kingsford, C. and Salzberg, S.L. (2008) What are decision trees? Nature biotechnology, 26, 1011–1013.

33. Breiman, L. (2001) Random forests. Machine learning, 45, 5–32.

34. Chen, C., Liaw, A. and Breiman, L. (2004) Using random forest to learn imbalanced data. *University of California*, Berkeley, 110, 24.

35. Webb, G.I., Keogh, E. and Miikkulainen, R. (2010) Naïve Bayes. Encyclopedia of machine learning, 15, 713–714.

36. Balakrishnama, S. and Ganapathiraju, A. (1998) Linear discriminant analysis-a brief tutorial. Institute for Signal and information Processing, 18, 1–8.

37. Liu, X.-Y., Wu, J. and Zhou, Z.-H. (2008) Exploratory undersampling for class-imbalance learning. *IEEE Transactions on Systems, Man, and Cybernetics*, Part B (Cybernetics), 39, 539–550.

38. Maclin, R. and Opitz, D. (1997) An empirical evaluation of bagging and boosting. AAAI/IAAI, 1997, 546–551.

39. Hido, S., Kashima, H. and Takahashi, Y. (2009) Roughly balanced bagging for imbalanced data. Statistical Analysis and Data Mining: The ASA Data Science Journal, 2, 412–426.

40. Seiffert, C., Khoshgoftaar, T.M., Van Hulse, J. and Napolitano, A. (2009) RUSBoost: A hybrid approach to alleviating class imbalance. IEEE transactions on systems, man, and cybernetics-part A: systems and humans, 40, 185–197.

41. Chen, T. and Guestrin, C. (2016), Proceedings of the 22nd acm sigkdd international conference on knowledge discovery and data mining, pp. 785–794.

42. Su, X., Yan, X. and Tsai, C.L. (2012) Linear regression. Wiley Interdisciplinary Reviews: Computational Statistics, 4, 275–294.

43. McDonald, G.C. (2009) Ridge regression. Wiley Interdisciplinary Reviews: Computational Statistics, 1, 93–100.

44. Tibshirani, R. (1996) Regression shrinkage and selection via the lasso. Journal of the Royal Statistical Society Series B: Statistical Methodology, 58, 267–288.

45. Zou, H. and Hastie, T. (2005) Regularization and variable selection via the elastic net. Journal of the Royal Statistical Society Series B: Statistical Methodology, 67, 301–320.

46. Awad, M., Khanna, R., Awad, M. and Khanna, R. (2015) Support vector regression. Efficient learning machines: Theories, concepts, and applications for engineers and system designers, 67–80.

47. Wang, J., Chen, Q. and Chen, Y. (2004), International symposium on neural networks. Springer, pp. 512-517.

48. Altman, N.S. (1992) An introduction to kernel and nearest-neighbor nonparametric regression. The American Statistician, 46, 175–185.

49. Freund, Y. and Schapire, R.E. (1997) A decision-theoretic generalization of on-line learning and an application to boosting. Journal of computer and system sciences, 55, 119–139.

50. Friedman, J.H. (2001) Greedy function approximation: a gradient boosting machine. Annals of statistics, 1189-1232.

51. Fischler, M.A. and Bolles, R.C. (1981) Random sample consensus: a paradigm for model fitting with applications to image analysis and automated cartography. Communications of the ACM, 24, 381–395.

52. Pedregosa, F., Varoquaux, G., Gramfort, A., Michel, V., Thirion, B., Grisel, O., Blondel, M., Prettenhofer, P., Weiss, R. and Dubourg, V. (2011) Scikit-learn: Machine learning in Python. the Journal of machine Learning research, 12, 2825–2830.

53. Lemaître, G., Nogueira, F. and Aridas, C.K. (2017) Imbalanced-learn: A python toolbox to tackle the curse of imbalanced datasets in machine learning. The Journal of Machine Learning Research, 18, 559–563.

54. Wright, E.P., Abdelhamid, M.A., Ehiabor, M.O., Grigg, M.C., Irving, K., Smith, N.M. and Waller, Z.A.E. (2020) Epigenetic modification of cytosines fine tunes the stability of i-motif DNA. Nucleic Acids Research, 48, 55–62.

55. Fojtík, P. and Vorlícková, M. (2001) The fragile X chromosome (GCC) repeat folds into a DNA tetraplex at neutral pH. Nucleic Acids Research, 29, 4684–4690.

56. Fleming, A.M., Ding, Y., Rogers, R.A., Zhu, J., Zhu, J., Burton, A.D., Carlisle, C.B. and Burrows, C.J. (2017) 4 n–1 is a “sweet spot” in DNA i-motif folding of 2′-deoxycytidine homopolymers. Journal of the American Chemical Society, 139, 4682–4689.

57. Brazier, J.A., Shah, A. and Brown, G.D. (2012) I-motif formation in gene promoters: unusually stable formation in sequences complementary to known G-quadruplexes. Chemical Communications, 48, 10739–10741.

58. Mir, B., Serrano, I., Buitrago, D., Orozco, M., Escaja, N. and González, C. (2017) Prevalent sequences in the human genome can form mini i-motif structures at physiological pH. Journal of the American Chemical Society, 139, 13985–13988.

59. Abdelhamid, M.A. and Waller, Z.A. (2020) Tricky topology: persistence of folded human telomeric i-motif DNA at ambient temperature and neutral pH. Frontiers in Chemistry, 8, 40.

60. Martella, M., Pichiorri, F., Chikhale, R.V., Abdelhamid, M.A., Waller, Z.A. and Smith, S.S. (2022) i-Motif formation and spontaneous deletions in human cells. Nucleic Acids Research, 50, 3445–3455.

61. Brooks, T.A., Kendrick, S. and Hurley, L. (2010) Making sense of G-quadruplex and i-motif functions in oncogene promoters. The FEBS journal, 277, 3459–3469.

62. Abou Assi, H., Garavís, M., González, C. and Damha, M.J. (2018) i-Motif DNA: structural features and significance to cell biology. Nucleic acids research, 46, 8038–8056.

63. Yazdani, K., Seshadri, S., Tillo, D., Yang, M., Sibley, C.D., Vinson, C. and Schneekloth Jr, J.S. (2023) Decoding complexity in biomolecular recognition of DNA i-motifs with microarrays. *Nucleic Acids Research*, gkad981.

